# Cold-induced skin darkening does not protect amphibian larvae from UV-associated DNA damage

**DOI:** 10.1101/2023.09.20.558735

**Authors:** Coen Hird, Emer Flanagan, Craig E. Franklin, Rebecca L. Cramp

**Author notes:** **Corresponding author:** Coen Hird.

## Abstract

Many amphibian declines are correlated with increasing levels of ultraviolet radiation (UVR). While disease is often implicated in declines, environmental factors such as temperature and UVR play an important role in disease epidemiology.
The mutagenic effects of UVR exposure on amphibians are worse at low temperatures. Amphibians from cold environments may be more susceptible to increasing UVR. However, larvae of some species demonstrate cold acclimation, reducing UV-induced DNA damage at low temperatures. Understanding of the mechanisms underpinning this response is lacking.
We reared *Limnodynastes peronii* larvae in cool (15°C) or warm (25°C) waters before acutely exposing them to 1.5 h of high intensity (80 μW cm^-2^) UVBR. We measured the colour of larvae and mRNA levels of a DNA repair enzyme. We reared larvae at 25°C in black or white containers to elicit a skin colour response, and then measured DNA damage levels in the skin and remaining carcass following UVBR exposure.
Cold acclimated larvae were darker and displayed lower levels of DNA damage than warm-acclimated larvae. There was no difference in CPD-photolyase mRNA levels between cold- and warm-acclimated larvae. Skin darkening in larvae did not reduce larval accumulation of DNA damage following UVR exposure.
Our results showed that skin darkening alone does not explain cold-induced reductions in UV-associated DNA damage in *L. peronii* larvae. Beneficial cold-acclimation is more likely underpinned by increased CPD-photolyase abundance and/or increased photolyase activity at low temperatures.

**Research Highlights:**

- *L. peroniii* larvae darken when exposed to cold temperatures
- Darker larvae were not protected from the effects of UV on DNA damage
- Cold acclimation of larvae when exposed to UV is likely driven by DNA repair enzymes not melanin

## 1. Introduction

We are living through the ‘sixth mass extinction’ event, and amphibians continue to approach extinction at alarming rates (McCallum, 2007). Many population declines initially classed as enigmatic were recorded from ‘pristine’ habitats, particularly at high altitudes associated with colder temperatures and higher ultraviolet radiation (UVR) levels. These declines are now largely linked to several complex and interacting factors, primarily disease and environmental change (Kiesecker et al., 2001; Carey and Alexander, 2003; Blaustein and Bancroft, 2007; Sodhi et al., 2008; Lötters et al., 2009; Alton and Franklin, 2017; Cramp and Franklin, 2018). Many amphibian larvae are sensitive to changes in temperature and UVR (Brattstrom, 1979; Blaustein et al., 1999, 2001, 2003; van Uitregt et al., 2007; Bancroft et al., 2008; Sapsford et al., 2013; Richter-Boix et al., 2015; Morison et al., 2019; Hird et al., 2022). The spatiotemporal correlation between global amphibian population declines and ozone depletion, as well as mounting evidence that climate change continues to influence UVR levels in aquatic habitats, has spurred decades of research into the detrimental effects of ultraviolet radiation (UVR) on amphibians (Carey, 1993; Barnes et al., 2019, 2022; Bornman et al., 2019; McKenzie et al., 2020). UVR exposure causes a range of mutagenic and cytotoxic effects that have negative outcomes for amphibians, primarily by damaging nucleic acids directly through the formation of photoproducts (primarily cyclobutene pyrimidine dimers [CPDs]) or indirectly through UV-induced oxidative stress and photosensitising structures (Lesser et al., 2001; Premi et al., 2015; Solano, 2016). Importantly, amphibians can repair UV-induced DNA damage through the action of DNA repair enzymes such as CPD-photolyase (Liu et al., 2015).

Although many amphibian declines have occurred at high altitudes associated with increased UVR and decreased temperatures, the interactive effects of UVR and temperature on amphibian health remain poorly understood. Lower temperatures compound the negative effects of UVR on amphibian embryos (Grant and Licht, 1995) and larvae (Broomhall et al., 2000). Larvae of the striped marsh frog (*Limnodynastes peronii*) experienced increased mortality, reduced growth and reduced swimming performance when exposed to high UVR doses at cooler temperatures (van Uitregt et al., 2007). Reduced growth results in smaller larvae, making individuals more susceptible to UVR-induced DNA damage due to their greater surface area to volume ratio. Cold temperatures have also been shown to delay metamorphosis, which extends the duration of UVR exposure (Garcia et al., 2003). DNA photorepair is thermally sensitive in amphibians and other ectotherms (Pakker et al., 2000; Li et al., 2002; MacFadyen et al., 2004; Lamare et al., 2006; Morison et al., 2019). *L. peronii* larvae acutely exposed to UVR at cold temperatures had increased levels of DNA damage in the form of CPDs compared to larvae exposed to UVR at warm temperatures (Morison et al., 2019). Other amphibian species have since demonstrated elevated DNA damage at cold temperatures (Hird et al., 2022), further suggesting that amphibians living in cold environments may be at elevated risk from increasing UVR levels.

Recently, however, *L. peronii* larvae reared and exposed to UVBR at cool temperatures accumulated less DNA damage than larvae reared in warm temperatures and acutely exposed to UVBR at cool temperatures (Hird et al., 2023a). This suggests that thermal acclimation, which is defined as a phenotypic change in response to the thermal environment within the lifetime of an organism (Kingsolver and Huey, 1998; Huey et al., 1999; Angilletta et al., 2006) can improve the resilience of larvae to UVBR-associated DNA damage. The degree to which amphibians demonstrate plasticity to changing abiotic factors in nature likely reflects their ability to cope with rapid global change (Kingsolver and Huey, 1998; Huey et al., 1999; Woods and Harrison, 2002; Angilletta, 2009; Chevin et al., 2010; Seebacher et al., 2014). The finding that thermal acclimation reduced UVR-induced DNA damage in L. peronii larvae at low temperatures suggests that aquatic ectotherms living in cool temperatures may be more resilient to high UVR than previously thought. However, the specific mechanisms underpinning this acclimation response are unknown.

Many adult amphibians are unable to thermally acclimate rates of physiological processes (Putnam and Bennett, 1981; Rome, 1983; Knowles and Weigl, 1990; Wilson and Franklin, 2000; Niehaus et al., 2011; Renaud and Stevens, 2011; Whitehead et al., 2015), but larval amphibians have a greater capacity for acclimation (Wilson and Franklin, 1999; Vo and Gridi-Papp, 2017). Cold acclimation in amphibian larvae leading to a decrease in UV-induced CPDs could be achieved through multiple physiological pathways. For example, thermal acclimation of CPD-photolyase activity or abundance could increase the efficacy of DNA damage repair at cold temperatures, offsetting the depressive effects of temperature on DNA damage accumulation.

Another way *L. peronii* larvae could reduce UV-induced DNA damage accumulation at cold temperatures is via the mobilisation of melanin to the epithelium, providing photoprotection during UVR exposure (Agar and Young, 2005; Franco-Belussi et al., 2016; Rodríguez-Rodríguez et al., 2020). *L. peronii* larvae turned visibly darker when reared in colder waters, possibly explaining why cold-acclimated larvae had less CPDs than warm-acclimated larvae when exposed to UVR, regardless of exposure temperature (Hird et al., 2023a). Although tadpole melanophores are mostly underneath the epidermis (Fox, 1986), melanin is often implicated in photoprotection from UVR (Kollias et al., 1991; Grant and Licht, 1995; Hofer and Mokri, 2000). However, many non-mutually exclusive hypotheses exist for the role of melanism in ectotherms such as aposematism, camouflage and ‘thermal melanism’ (Clusella Trullas et al., 2007).

The thermal melanism hypothesis posits that dark ectotherms are at a thermal advantage compared to light ectotherms at cold temperatures because they can heat up faster (Rodríguez-Rodríguez et al., 2020). However, caution is needed when suggesting that darker colouration causes thermoregulatory advantages (Clusella Trullas et al., 2007). Thermal melanism is hard to explain in small aquatic ectotherms due to effective convective heat transfer in water as a consequence of aquatic respiration, and the high thermal conductivity and specific heat capacity of water compared with air (Grigg et al., 1979; Manning and Grigg, 1997; Haesemeyer, 2020). This makes it theoretically impossible for small aquatic ectotherms to regulate their body temperature apart from behavioural thermoregulation within the water column (but see Nordahl et al., 2018). Therefore, the cold-induced darkening of amphibian larvae observed by Hird et al. (2023) was unlikely to be thermoregulatory and may instead play a role in photoprotection at lower temperatures. The dispersal of melanin throughout the dermal layer, resulting in a darker colouration, has been shown to protect against UVR-induced DNA damage by reducing the depth at which UVR wavelengths can penetrate the body (Kollias et al., 1991; Hofer and Mokri, 2000; Agar and Young, 2005). Dermal melanin absorbs UVR, resulting in a UVR screening effect which may localise UV-induced DNA damage to the skin. We propose a beneficial thermal acclimation hypothesis to explain cold-induced darkening in amphibian larvae, where animals darken at cool temperatures to compensate for the depressive effects of temperature on UV-induced DNA damage.

In this study, we explored the mechanistic underpinnings for the reduction in DNA damage levels in cold-acclimated *L. peronii* larvae observed by Hird et al. (2023) to test two potential mechanisms of thermal acclimation. To test whether cold-acclimation either induced cutaneous darkening and/or promoted the upregulation of CPD-photolyase expression, *L. peronii* larvae were chronically exposed to cool (15°C) or warm (25°C) temperatures for three weeks and larval colour and CPD-photolyase mRNA levels were quantified. It was hypothesised that larvae would acclimate to cold temperatures by upregulating CPD-photolyase mRNA and mobilising melanin to the epithelium (skin darkening), both serving to reduce potential DNA damage from UVR. To determine whether skin darkening alone is a photoprotective mechanism, in the context of reducing UV-induced CPDs, *L. peronii* larvae were reared in a common environment and acclimated to light or dark backgrounds to stimulate skin darkening. It was hypothesised that acclimation to dark backgrounds would cause rapid skin darkening comparable to cold acclimation, which would result in reduced DNA damage following an equal UVR exposure compared with larvae acclimated to a light background.

## 2. Materials and methods

### 2.1. Experimental animals and general methods

Six part clutches of freshly laid *Limnodynastes peronii* spawn were collected near Magandjin Brisbane on Yuggera Country in Summer 2022 (-27.630595, 152.370432; QLD, Australia) and immediately transported to The University of Queensland. Spawn were transported to a mesocosm (PTT12W, Rapid Plas, Tamworth, NSW, Australia) filled with 150 L rainwater and covered with 70% UV-blocking white shade cloth. pH was monitored every second day (LAQUA P-22, Horiba Instruments, Singapore). Temperature was logged using ibuttons (type DS1921H; Maxim/Dallas Semiconductor Corp., USA; Figure S1) until larvae reached Gosner stage 25 (Gosner, 1960). N = 120 larvae were randomly moved from the mesocosms to the laboratory environment and split across three experiments in two-litre ice-cream containers half-filled with carbon-filtered Brisbane City tap water. Larvae were fed frozen spinach ad libitum and partial water changes were conducted every 2-3 days. Larvae were reared under a 12L:12D photoperiod generated using standard roof fluorescent lights. UVR for experimental exposures was generated using 40 W, full spectrum fluorescent lights which emit visible light, UV-A, and UV-B (Repti-Glo 10.0, 1200 mm, Exo Terra, Montreal, Canada). Light intensities for UVB were measured using a radiometer (IL1400BL, International Light Inc., Newburyport, USA; Table 1).

**Table 1.**
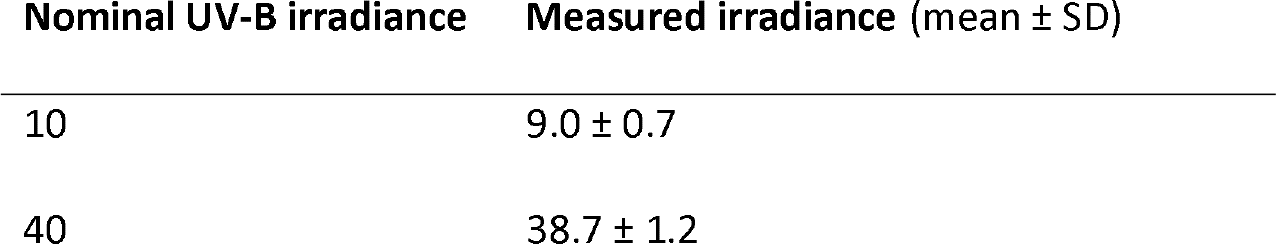
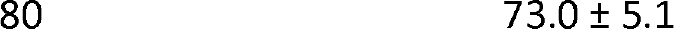
Measured UV-B values for each UVR chronic and acute treatment. Chronic (10 and 40 μW cm^-2^ treatments were exposed for 1 h daily. Acute (80 μW cm^-2^) UVR exposure was for 1.5 h in Experiments 1 and 2, and for 1 h in Experiment 3. All values are reported in μW cm^-2^.

### 2.2. Experimental design

#### 2.2.1. Experiment 1

Spawn (N = 40) were reared in individual 1 L clear plastic containers (BBC Plastics, Sydney, Australia). This experiment used the same experimental design as Hird et al. (2023) since it sought to understand the mechanistic basis for the improved photoprotection of cold-acclimated *L. peronii* larvae documented in that study. Water temperatures were achieved by holding spawn in separate temperature-controlled rooms where they were maintained for the duration of the experiment. Larvae were held at 25°C or 15°C at either a low or moderate UV-B lighting treatment (in a 2x2 factorial design) which corresponded to approximately 10 or 40 μW cm^-2^ at the water surface administered for 1 h daily (at midday).

Larvae were maintained under their treatment conditions for 21 days post-hatching. Larvae were individually placed into separate wells in six-well plates containing 10 ml of Brisbane carbon filtered tap water. The plates were evenly distributed across two water baths (volume 32 L; L 650 mm x W 410 mm x H 210 mm; water depth ∼160 mm) at 15°C or 25°C. The temperature of the experimental water baths was controlled using 300 W aquarium heaters (AquaOne) and water was circulated by small aquarium pumps to ensure thermal uniformity and to move plates across the surface of the water to homogenise UVR received by larvae. Larvae were left for one hour to adjust to the experimental temperature. Prior to and immediately following UVR exposure, photos were taken of larvae from a fixed height using a Canon EOS 90D camera with the following manual settings: 1/125 shutter speed; F8.0 aperture; ISO 3200, shot in JPG file format. Photos were taken of larvae at a consistent water depth, using an identical grey background with scale and diffused lighting to wholly illuminate each tadpole. UVR lights were switched on at 12:00pm and all larvae were exposed to ∼80 μW cm^-2^ UV-B for 1.5 hours (table 1). UV lights were then covered in UVB blocking film (Melinex 516, 100 um, Archival Survival, Doncaster, VIC, Australia) and larvae were left to photorepair damaged DNA for 12 h, representing the time point at which maximal CPD-photolyase mRNA levels occurred following UV-induced upregulation (Morison et al., 2019). Afterwards, larvae were euthanised in buffered MS222 (0.25 mg L^-1^; Ramlochansingh et al., 2014) and stored in RNAlater (Life Technologies, Carlsbad, CA, USA) at 4°C for 24 h, before being moved to -20°C.

#### 2.2.2. Experiment 2

At 100 days post-hatch, larvae held at 25°C (n = 40) were moved to 950 ml black (n = 20) or white (n = 20) L noodle bowls (CONPLNB950B, CONPLNB950WHI, Statpack, Rocklea, QLD, Australia) for 24 h to allow them time to adjust their body colour to match their backgrounds (Rodríguez-Rodríguez et al., 2020). Larvae were then acutely exposed to 80 μW cm^-2^ UV-B for 1.5 hours at 25°C housed in six-well plates as in experiment 1. All subsequent steps were conducted under red light to prevent photoenzymatic repair (PER) occurring after the experiment. Prior to and immediately following UVR exposure, photos were taken of larvae as in experiment 1. Larvae were euthanised by immersion in MS222 and snap frozen in dry ice and stored at -80°C.

#### 2.2.3. Experiment 3

Larvae used in experiment 3 (n=40) were maintained at 23°C. At day 157 post-hatch, larvae were transferred to individual black (n=5) or white (n=5) bowls. After 24 hours, larvae were transferred into individual wells in 6-well plates containing 10 ml of filtered tap water. Larvae were photographed immediately prior to UVR exposure from a fixed height using a Google Pixel 6 phone camera. Photos were taken of larvae at a consistent water depth, using an identical background with a scale and diffused lighting to wholly illuminate each tadpole. Larvae were then acutely exposed to UVR as in experiment 1 and 2. All subsequent steps were conducted under red light to prevent PER occurring after the experiment. Larvae were then euthanised by immersion in 250 mg L^-1^ benzocaine solution. Tadpole skin was removed from the carcass, and both were snap frozen in dry ice, and stored at -80°C for subsequent CPD detection.

### 2.3. Image analyses

Skin colour was assessed using ImageJ (National Institutes of Health, Bethesda, Maryland, USA). Images were converted to 8-bit greyscale. Pictures were normalised by calculating pixel values of the image sequence, so the range is equal to the maximum range for the data type, using an in-built normalisation function. An average reading of pixel values ranging from 0 (pure black) to 255 (pure white) was taken over a specified portion of the dorsal surface on each tadpole, just behind the eyes of the tadpole (Lundsgaard et al., 2021).

### 2.4. CPD detection

Genomic DNA was extracted and purified from whole-larvae, skin, or carcass homogenates using a Qiagen DNeasy Blood and Tissue Kit (Qiagen Inc., Hilden, Germany) and quantified using a Qubit dsDNA High-Sensitivity Assay Kit (ThermoFisher Scientific Inc., Waltham, MA, USA). CPD concentrations were determined using an anti-CPD ELISA assay following the primary antibody manufacturer’s protocol (Mori et al., 1991; NM-ND-D001, clone TDM-2, Cosmo Bio Co., Ltd.) and following Hird et al. (2023). Plates were read at 450 nm in a DTX880 multimode detector (Beckman Coulter, MN, USA) using the SoftMax® Pro program (Version 7.1.0, Molecular Devices LLC, CA, USA). CPD concentrations were calculated from a standard dose-response curve of UVC-irradiated calf thymus (NM-MA-R010, Cosmo Bio Co., Ltd.) present on each plate. CPD concentration is reported as units of UVCR-dose equivalents per 20 ng of DNA.

### 2.5. CPD-photolyase gene expression

Total RNA was extracted from *L. peronii* using a RNeasy Plus Mini Kit following the manufacturer’s instructions (Qiagen, Valencia, CA, USA) and including the gDNA Eliminator Spin Column step. Total RNA was eluted from the silicon spin column in elution buffer and its concentration quantified using a Qubit fluorometer (ThermoFisher Scientific, Waltham, MA, USA). Residual genomic DNA contamination was removed using DNase I (ThermoFisher Scientific, Waltham, MA, USA) and RNA was then reverse transcribed using an iScript cDNA Synthesis Kit (Bio-Rad Laboratories Inc., Hercules, CA, USA) following the manufacturers guidelines. Appropriate no-reverse transcriptase controls were generated by replacing reverse transcriptase with water. No-reverse transcriptase controls were assessed independently for each biological sample to confirm the absence of genomic DNA contamination.

Primer sequences for *L. peronii* CPD-photolyase and the house-keeping gene beta actin were obtained from Morison et al. (2019). Primers were re-evaluated for specificity and to ensure that they produced only a single band of the appropriate length using MyTaq DNA Polymerase (Bioline, Alexandria, NSW, Australia) and agarose gel electrophoresis.

Quantitative PCR assays were conducted using iTaq™ Universal SYBR Green Supermix (Bio-Rad Laboratories Inc.) in a CFX Connect Real-Time PCR Detection System (Bio-Rad Laboratories Inc., REFS). Samples were analysed in triplicate in 20 ul reactions using neat cDNA, and each plate included appropriate no-template controls. Cycling parameters were as follows: 94°C for 2 min followed by 39 cycles of 94°C for 15 s and 60°C for 30 s. Melt (dissociation) curves (65-90°C) were conducted after each run. Reaction efficiencies were calculated using a serially diluted pooled cDNA standard curve. All PCR efficiencies were greater than 90% with an R^2^ of over 0.99. All assays produced unique and single peak dissociation curves. Data were exported to MS Excel using Bio-Rad CXF Manager (version 3.1). The following calculations for CPD-photolyase gene expression were conducted following Pfaffl (2001):

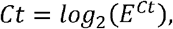

where *E* is the reaction efficiency for the gene and *Ct* is the raw *Ct* value of the sample. Δ*Ct* was calculated for each sample for statistical analyses:

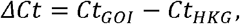

where *Ct*_*GOI*_ is the *Ct* value for the gene of interest, and *Ct*_*HKG*_ is the Ct value for the housekeeping gene. Δ*Ct* in each target gene was quantified as fold-change relative to the expression of a reference group (25°C and 10 μW cm^-2^ UV-B acclimated *L. peronii* larvae, n = 5).

### 2.6 Statistical analyses

All analyses were conducted in the R statistical environment (R Core Team, 2023). Regression diagnostics were analysed using the performance package (Lüdecke et al., 2021). Models were two-tailed, assumed a Gaussian error structure and all models met the assumptions of the statistical tests used. α was set at 0.05 for all tests (but see experiment 3 below). The effect of wet body mass on CPD accumulation was accounted for by fitting wet body mass as a covariate in the model.

Raw Ct values for the housekeeping gene were compared between treatments for experiment 1 in an analysis of variance model using the car package (Fox and Weisberg, 2018), to assess the stability of the gene across treatments. Differences in CPD-photolyase gene expression between treatment groups in experiment 2 were analysed by comparing Δ*Ct* values also in an analysis of variance model.

Raw colour data of larvae in experiment 1 and 2 were fitted in separate linear mixed effects models using tadpole ID as a random effect to account for repeated measurements. Colour differences in experiment 3 were fitted in in an ANOVA with background colour as the independent variable. Colour differences between treatment groups within the model for all experiments were compared using Type II Wald F tests with Kenward-Roger degrees of freedom in the car package (Fox and Weisberg, 2018). Contrasts between fixed effects and post hoc analyses were obtained using the sjPlot package (Lüdecke, 2021) and the emmeans package (Lenth, 2020). The influence of the random effects on the model was assessed using REML likelihood ratio tests with the package lmerTest (Kuznetsova et al., 2017).

DNA damage data from experiment 2 and 3 were modelled using one-way ANCOVAs incorporating mass as a covariate and acclimation background colour as an independent variable. Differences in CPD accumulation between treatment groups within the model were compared using Type II sums of squares with the car package (Fox and Weisberg, 2018). Alpha was set to 0.025 by Bonferroni correction for comparing DNA damage data from experiment 3 due to multiple univariate tests (skin and carcass). All estimates reported from models are estimated marginal means adjusted for the effect of the covariate. Any random effects of 6-well plate ID during UV exposure or housing on rearing were randomly distributed across the treatments and were not modelled as random effects due to issues with low group sizes in modelling and low likelihood of directly influencing the response variable. Due to slight differences in experimental design and data collection, no statistical comparisons were made between experiments for any trait. A copy of the script used to analyse these data are available at UQ eSpace (doi:10.48610/f559345).

## 3. Results

### 3.1. Experiment 1

Cold-acclimated larvae were smaller on average (n=20; mean ± SD: 26.5 mg ± 9.4) than warm-acclimated larvae (n=20; mean ± SD: 35.4 mg ± 13.7). Larvae reared at 15°C were darker than larvae reared at 25°C, though the magnitude of the effect was greater in larvae photographed prior to the acute UVR exposure (Figure 1A; F_1,73_ = 4.75, *p* < 0.03). There was no effect of UVR acclimation on colour change in *L. peronii* larvae. Analyses of ΔCt values showed no effect of UVR (F_1,15_ = 0.23, *p* = 0.64) or temperature (F_1,15_ = 0.08, *p* = 0.79) acclimation on CPD-photolyase mRNA expression levels following 24 hours of DNA photorepair (Figure S2).

**Figure 1.**
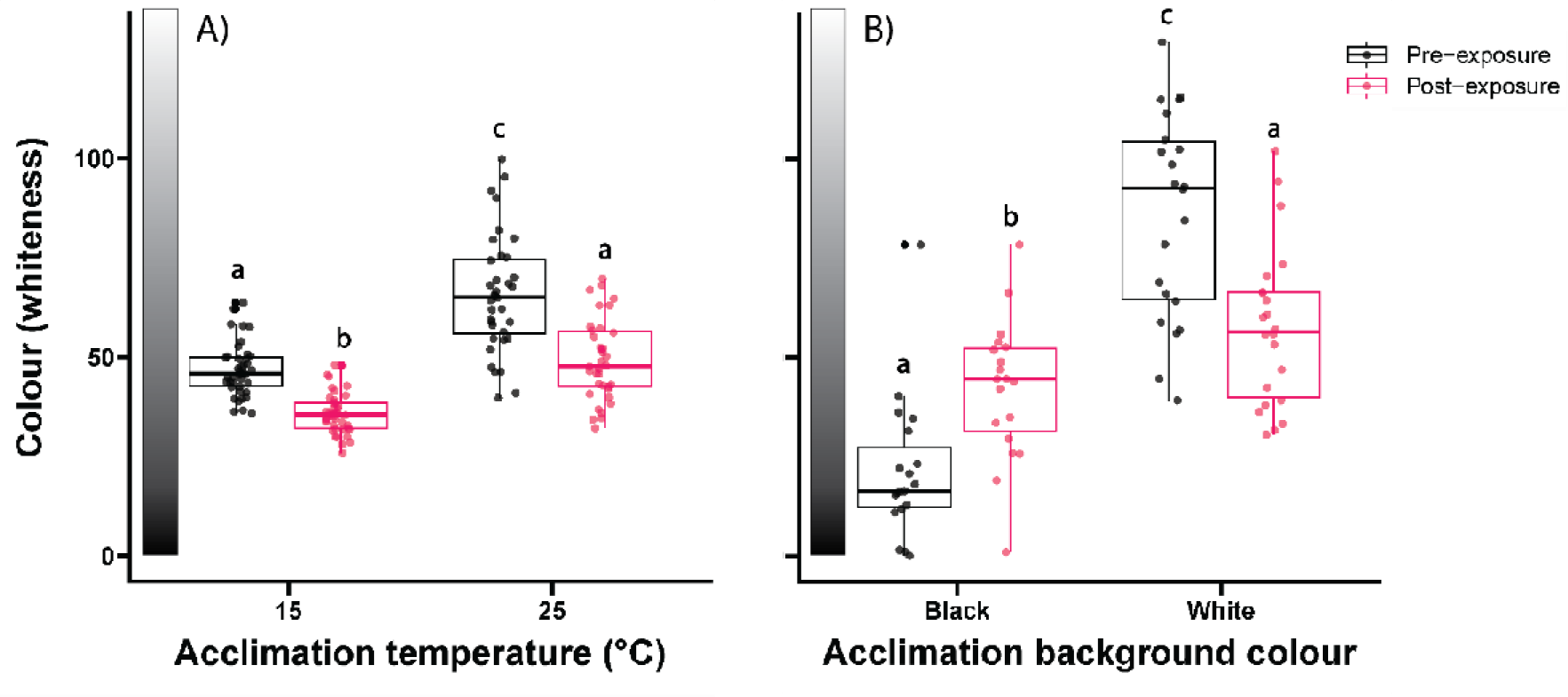
The effect of acclimation temperature on body colour in *L. peronii* larvae prior to and immediately following 1.5 hours of acute high UVR exposure (80 μw cm^-2^ UV-B) in A) larvae acclimated to cool (15°C) and warm (25°C) waters and B) larvae acclimated to black or white backgrounds. Lower colour values represent darker individuals. Small circles represent individual raw data points. Different lower-case letters represent statistically significant differences between groups.

3.2. Experiment 2

Larvae (n=40) were larger on average (mean ± SD: 59.1 mg ± 37.1) than larvae used in experiment 1. Larvae acclimated to dark backgrounds for 24 hours were significantly (∼300%) darker than larvae acclimated to white backgrounds, though this effect was only present in larvae photographed prior to the acute UVR exposure (Figure 1B; *t*_78_, F_*1,78*_ = 28.39, *p* <0.001). Following UVR exposure, dark-acclimated larval bodies were ∼100% lighter (*t*_39_, *p* <0.01) and light-acclimated larval bodies were ∼75% darker (*t*_69_, *p* = <0.001). There was weak evidence that larvae exposed to black backgrounds were still significantly darker than larvae exposed to white backgrounds following UVR exposure (*t*_69_, *p* = 0.09). There was no effect of acclimation to dark or light backgrounds on whole body DNA damage in the form of CPDs following acute UVR exposure (Figure S3; F_1,37_ = 1.71, *p* = 0.2).

### 3.3 Experiment 3

Larvae (n=40) were larger on average (mean ± SD: 73.8 mg ± 20.2) than larvae used in experiment 2 and 3. Larvae acclimated to dark backgrounds for 24 hours were significantly darker than larvae acclimated to white backgrounds (Figure S4; F_1,34_ = 180.97, *p* < 0.001). There was no effect of acclimation to dark or light backgrounds on DNA damage in the form of CPDs following acute UVR exposure in either the larval skin (F_1,34_ = 0.41, *p* = 0.53) or the carcass (F_1,34_ = 0.26, *p* = 0.62; Figure 2).

**Figure 2.**
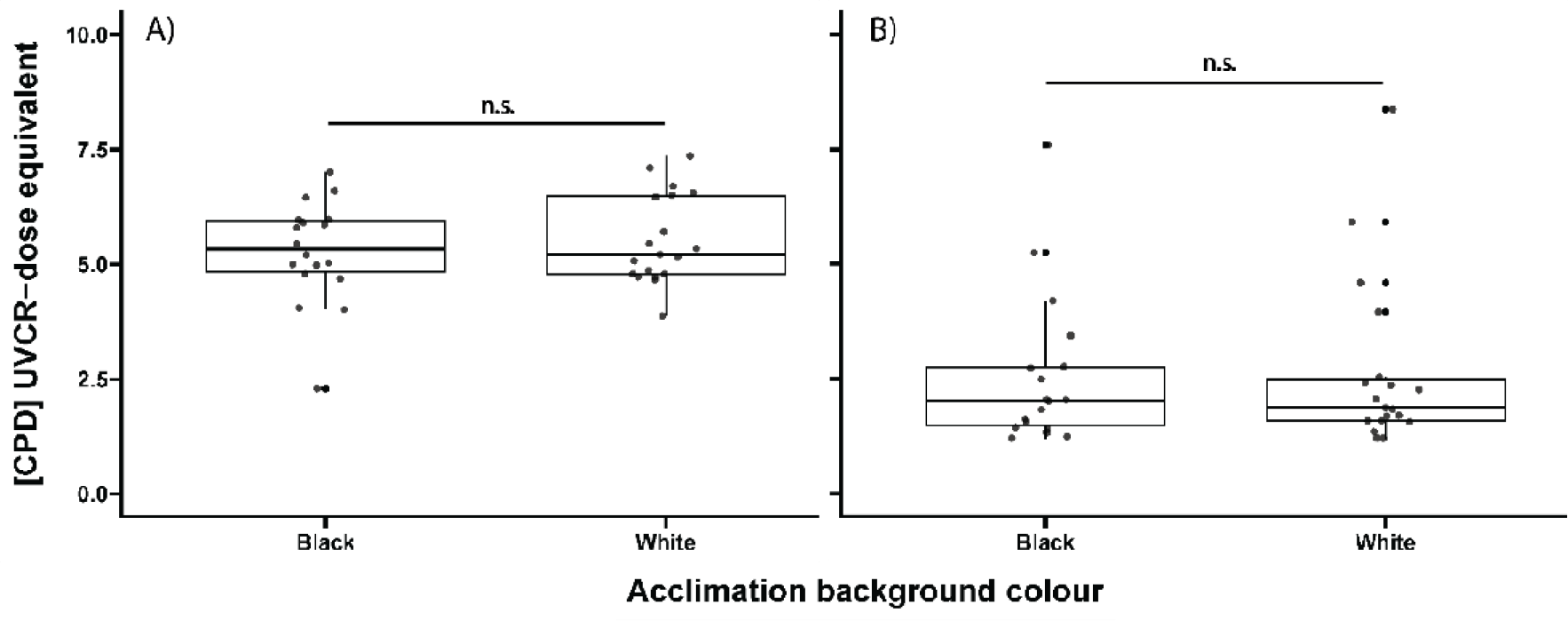
The effect of acclimation background colour on DNA damage concentration in the form of CPDs in *L. peronii* larval A) skin and B) the skin-free carcass prior to and immediately following 1.5 hours of acute UV-B exposure (80 μw cm^-2^). Small circles represent individual raw data points.

## 4. Discussion

*L. peronii* larvae acclimated to cooler temperatures elicited a darkening response, supporting some but not all studies on factors influencing melanin dispersion in amphibians (Norris and Milstead, 1967; Kats and van Dragt, 1986; Moriya and Miyashita, 1989; Fernandez and Bagnara, 1991; Jenks et al., 2007; Malik et al., 2023). Against our hypothesis, darker larvae acclimated to dark backgrounds did not have reduced levels of DNA damage in the form of whole-body or skin CPDs following UVR exposure compared to lighter larvae. This supports findings in newts and salamanders, where the presence of increased melanin also did not influence DNA damage (Lesser et al., 2001) or improve survival (Bancroft, 2007; Belden & Blaustein, 2002) following UVR exposure. These results suggested that skin darkening provided no photoprotection against direct UVR-induced DNA damage in *L. peronii* larvae. This result is especially surprising considering that eumelanin, the main pigment involved in the amphibian darkening response, typically provides photoprotection (Giuseppe Prota, 1992; Frost-Mason and Mason, 2004; Galván et al., 2011; Ito et al., 2018). Cold-induced darkening may be non-adaptive, related to another adaptive purpose, or its photoprotection was undetectable due to reduced DNA repair rates caused by cool temperatures.

Contrary to our hypothesis, acclimation to cold temperatures was not associated with changes to CPD-photolyase mRNA expression levels at 12 hours post UVR-exposure in *L. peronii*. This period was associated with the highest levels of CPD-photolyase upregulation following UVR exposure in an earlier study of *L. peronii* larvae (Morison et al. 2019). However, it is possible that cold-acclimation could induce the upregulation of CPD-photolyase following UVR exposure earlier than in warm-acclimated animals. To our knowledge there is no evidence of changes in the expression of DNA photorepair genes following thermal acclimation in animals, however acclimation to progressively colder temperatures significantly increased the upregulation of photolyase expression in wheatgrass (Jaikumar et al., 2020). While not direct evidence of an increase in photolyase gene expression, *Drosophila* selectively bred to cooler temperatures demonstrated upregulation of enzymes associated with nucleotide excision repair (Telonis-Scott et al., 2009). (Jaikumar et al., 2020).

Organisms acclimated to cool temperatures may increase constitutive photolyase concentrations to compensate for the depressive effects of low temperatures on DNA damage repair rates. Our findings suggested that reduced UV-induced DNA damage associated with cold-acclimation in *L. peronii* larvae (Hird et al., 2023a) may be caused by other mechanisms. To determine this, photolyase gene expression levels before and following UVR exposure in cold-acclimated larvae are needed. Amphibian larvae may also have capacity for thermal acclimation of photolyase activity (suggested by Morison et al. 2019), increasing the rate of DNA repair following UVR exposure at low temperatures. Investigating photolyase activity and abundance over time prior to and in the hours following UVR exposure in cold-acclimated versus warm-acclimated amphibians is required to explain whether increases in CPD-photolyase abundance/activity underpin *L. peronii* cold-acclimation reducing UVR-induced DNA damage (Hird et al., 2023a).

## 5. Conclusions

Understanding the consequences of current rapid climate change for the natural world is a “grand challenge in ecology” (Thuiller, 2007). In ectothermic animals, the capacity to respond to environmental variability in temperature by thermal acclimation is well established (Lagerspetz, 2006). With mounting evidence that climate change is influencing UVR and thermal regimes in aquatic ecosystems (Sokolova and Lannig, 2008; Barnes et al., 2019, 2022; Bornman et al., 2019; McKenzie et al., 2020), as well as species’ altitudinal distributions and phenological shifts (Walther et al., 2002; Bush et al., 2007; Hoffmann and Sgró, 2011; Cohen et al., 2018; Chmura et al., 2019), many aquatic ectotherms are and will be exposed to new combinations of UVR and temperature. Considering the susceptibility of aquatic ectotherms to UVR at cold temperatures, it is important to understand how physiological mechanisms explain their capacity for acclimation and ultimately tolerance of UVR exposure at different temperatures. It is difficult to tease apart the relative contributions of different physiological mechanisms that underpin the UVR-induced DNA damage response. However, our results strongly suggest that skin darkening is not enough to explain cold-induced reductions in UV-induced DNA damage in *L. peronii* larvae. This beneficial cold-acclimation is more likely underpinned by increased constitutive CPD-photolyase abundance and/or activity at low temperatures.

## Acknowledgements

We acknowledge Yuggera Country where the animals in this research were collected and acknowledge the significance of amphibians to Indigenous cultures globally, recognising the need for appreciation of the cultural significance of more-than-human kin in experimental biology (Hird et al., 2023b). We thank the volunteers who assisted with animal maintenance for this project.

## Data availability

The complete datasets and R scripts used for analysing the data are publicly available at UQ eSpace (doi:10.48610/f559345).

## Author contributions

Conceptualisation: RLC, CH, CEF; Methodology: CH, RLC, CEF; Validation: CH; Formal analysis: CH; Investigation: CH; Resources: CEF; Data curation: CH; Writing – original draft: CH; Writing – review & editing: CH, RLC, CEF; Visualisation: CH; Supervision: RLC, CEF; Project administration: RLC, CEF; Funding acquisition: RLC, CEF.

