## Supplemental files for "Cold-induced skin darkening does not protect amphibian larvae from UV-associated DNA damage"

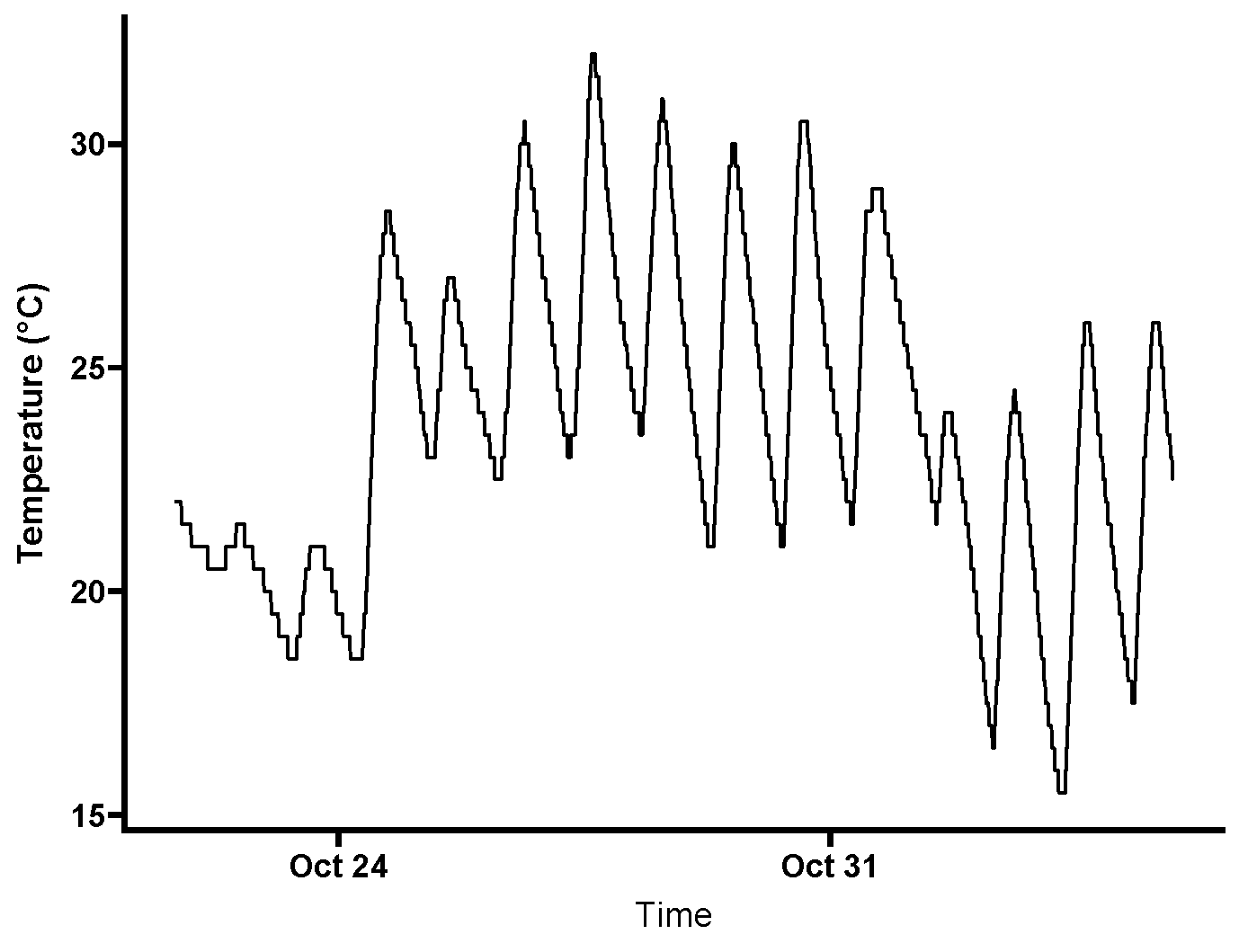

**Figure S1.** Water temperature measured in mesocosm *Limnodynastes peronii* rearing environment during development to Gosner stage 25.

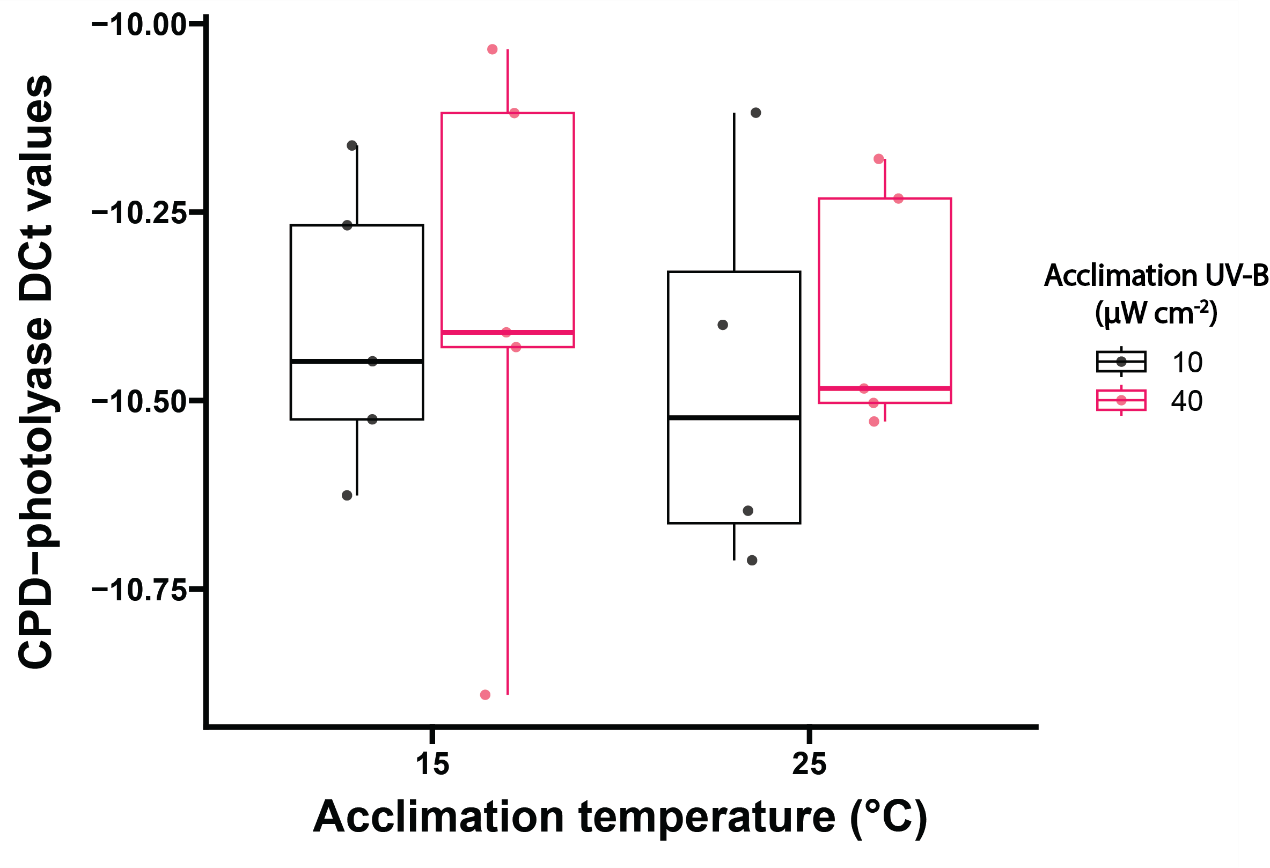

**Figure S2.** Effect of acclimation temperature and UV radiation on expression of CPD-photolyase in *L. peronii* larvae at 12 hours post UV-B exposure (1.5 h at 80 µW cm^-2^). Small circles represent delta Ct values for each larvae.

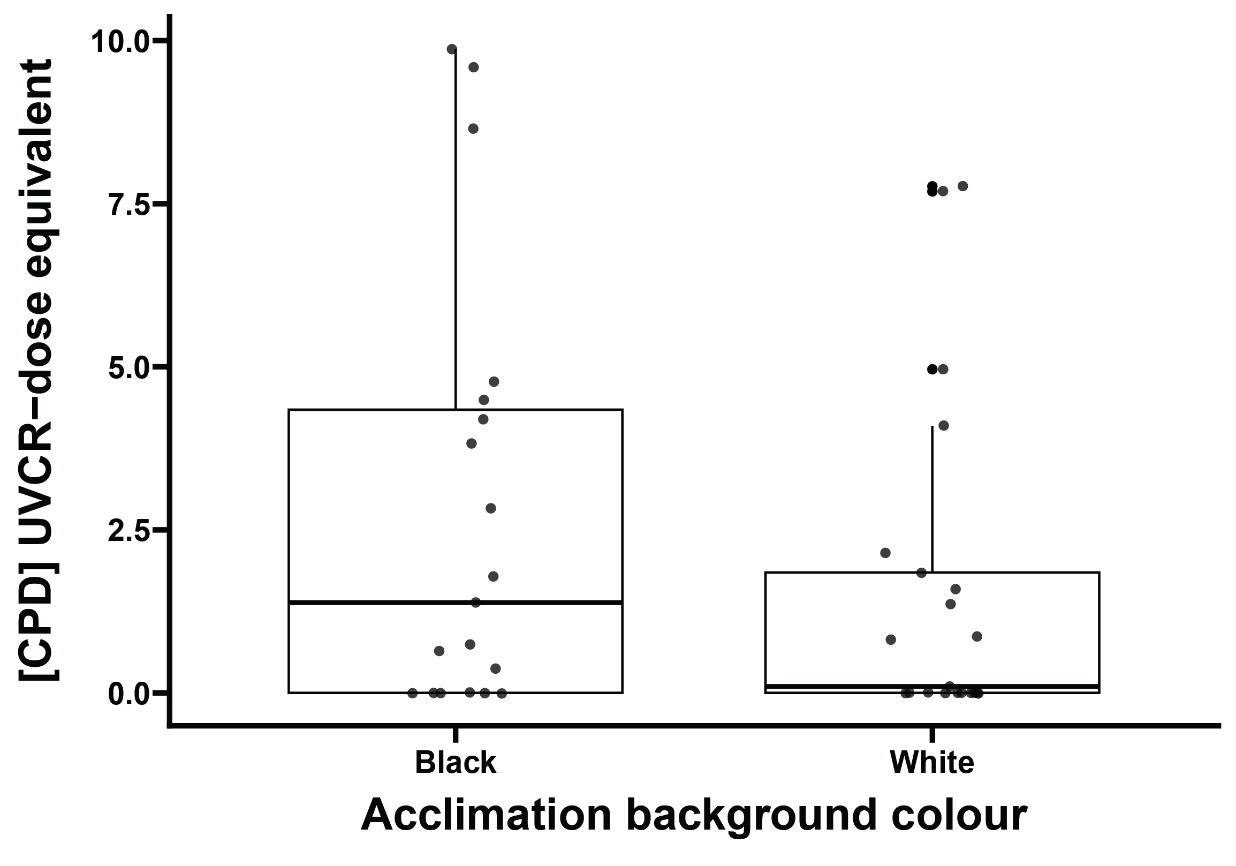

**Figure S3.** Effect of acclimation to dark or light backgrounds on DNA damage in form of CPDs in whole body *L. peronii* following an acute high UV exposure (1.5 h at 80 µW cm^-2^). Small circles represent CPD concentration of individual larvae.

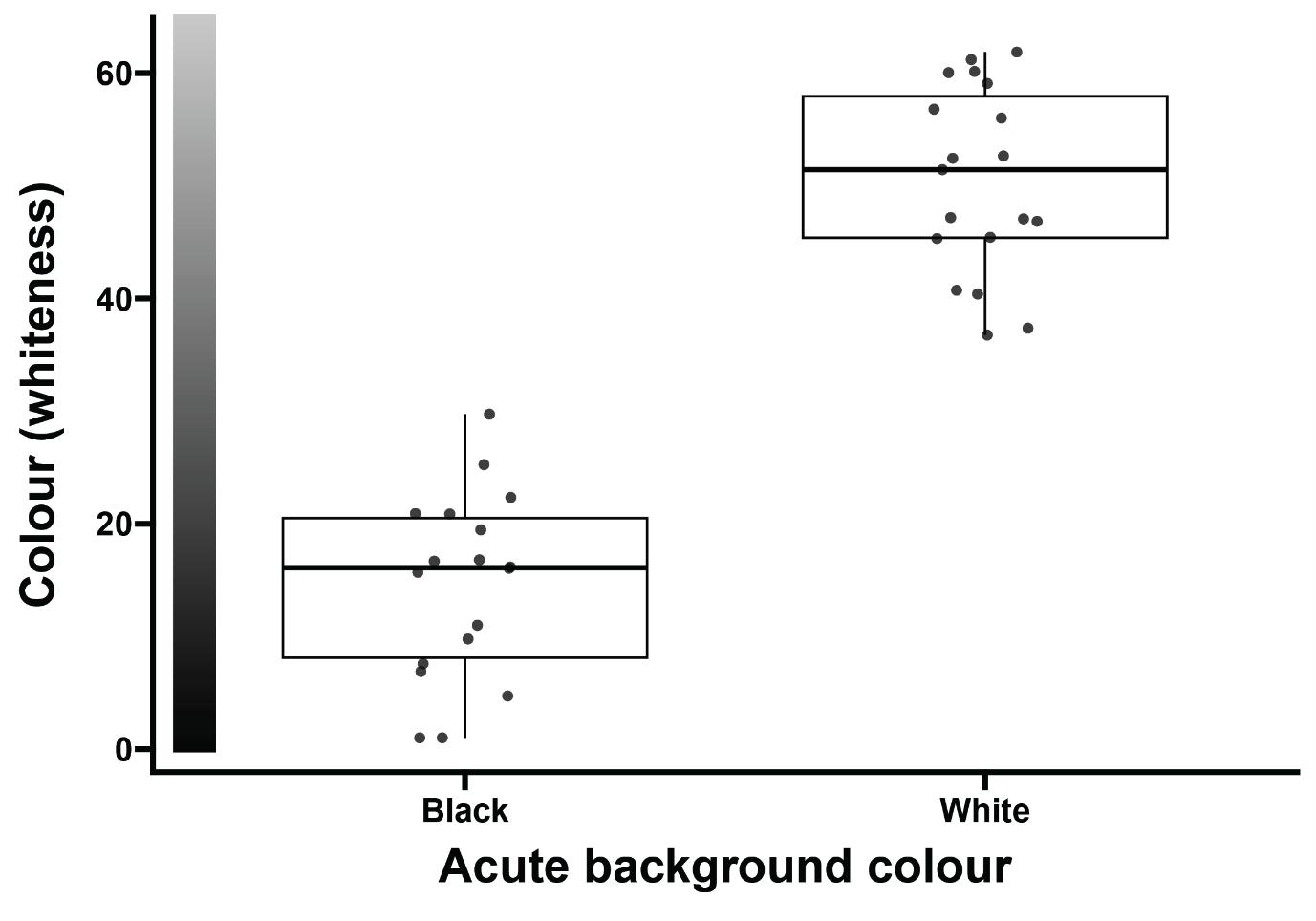

**Figure S4.** Effect of acclimation to dark or light backgrounds on larval body colour in *L. peronii* immediate prior to an acute high UV exposure (1.5 h at 80 µW cm^-2^). Small circles represent the colour of individual larvae. Lower values are associated with darker individuals.
